# Long-Term Effects of Early Adolescent Second-Generation Antipsychotic Exposure on Body Weight, Caloric Intake, and Metabolism-Associated Gene Expression

**DOI:** 10.1101/2024.09.29.615669

**Authors:** Paul L. Soto, Michael E. Young, Serena Nguyen, Megan Federoff, Mia Goodson, Christopher D. Morrison, Heidi M. Batdorf, Susan J. Burke, J. Jason Collier

**Affiliations:** Department of Psychology, Louisiana State University, Baton Rouge, LA 70803; Pennington Biomedical Research Center, Baton Rouge, LA 70808; Kansas State University, Manhattan, KS 66506

**Keywords:** second-generation antipsychotics, early adolescent exposure, weight gain, metabolic, mice, risperidone, olanzapine

## Abstract

The use of second-generation antipsychotic (SGA) medications in pediatric patients raises concerns about potential long-term adverse outcomes. The current study evaluated the long-term effects of treatment with risperidone or olanzapine on body weight, caloric intake, serum insulin, blood glucose, and metabolism-associated gene expression in C57Bl/6J female mice. Compared to mice treated with vehicle, female mice treated with risperidone or olanzapine gained weight at higher rates during treatment and maintained higher body weights for months following treatment cessation. During high-fat diet feeding, some groups of treated mice gained weight at higher rates than their respective control groups, but the finding was not consistent across experiments. Finally, female mice previously treated with olanzapine also exhibited increased expression of genes associated with inflammation and lipogenesis. These findings suggest that pediatric use of SGA medications that induce excess weight gain during treatment may exert persistent effects on body weight and gene expression and such outcomes may form an important aspect of assessing risk-to-benefit ratios in prescribing decisions.

---

Second-generation antipsychotic (SGA) medications are FDA-approved in the United States for use in pediatric patients as young as 5 years old (Christian et al., 2012; McNeil S.E. et al., 2023, Jan 16). In addition to their use for FDA-approved indications, prescribing data indicates that off-label usage of SGA medications in children and adolescents is common and may represent the vast majority of SGA prescriptions in children and adolescents (Olfson et al., 2012; Pathak et al., 2010). For example, in a review of outpatient prescribing records, Olfson et al. (2012) found that in children 3 years old and younger, 94% of office-based physician visits with antipsychotic treatment involved an off-label indication. Similarly, Olfson et al. (2012) found that in adolescents aged 14-20 years, 87% of office-based physician visits with antipsychotic treatment involved an off-label indication. Off-label usage of SGA medications is not limited to the United States. In a 2017 analysis of prescribing data from Australia, 79.7% of antipsychotic prescriptions in children and adolescents were off-label (Klau et al., 2024).

In general, use of psychotropic medications in children and adolescents raises concerns about potential biological and behavioral changes that may occur (Andersen & Navalta, 2004; Carlezon & Konradi, 2004; Steiner et al., 2014). SGA medications are specifically associated with above normal weight gain and metabolic and hormonal changes during treatment (Correll et al., 2009; De Hert, Dobbelaere, et al., 2011) and it remains unclear whether there are long-term risks to the use of SGA medications in pediatric populations, even after treatment discontinuation. Concerns about pediatric SGA medication usage have been noted because of the lack of data on long-term consequences (Bretler et al., 2019; Correll et al., 2013; Radojcic et al., 2023; Smith et al., 2022). Further, research on the long-term consequences of SGA medication use in pediatric populations has been identified as a high priority research area (Christian et al., 2015) and the need for assessments of the long-term safety of SGA medications has been noted by the American Academic of Child and Adolescent Psychiatry (American Academy of Child and Adolescent Psychiatry, 2011).

The current study evaluated the long-term effects of early adolescent exposure to SGA medication on body weight, caloric intake, gene expression, and circulating insulin and glucose in mice. We focused on body weight because above normal weight gain during childhood and adolescence is predictive of later adverse health outcomes (Balasundaram & Krishna, 2023; Barton, 2012) and although limited, data from studies in pediatric populations indicate that weight gain effects of SGA medications are, at best, only partially reversible (Doane et al., 2022; Speyer et al., 2021; Upadhyay et al., 2019).

We selected risperidone and olanzapine for these studies because of their risk of weight gain and their common use in pediatric populations. Risperidone and olanzapine have been classified as carrying intermediate and substantial risk of inducing weight gain, respectively (De Hert, Detraux, et al., 2011). As our goal was to evaluate whether body weight gains induced by medication treatment would persist following treatment cessation, we needed to use medications likely to produce above-normal weight gain. Although the prevalence of prescribing varies by patient group, insurance coverage, and likely many other variables, several recent retrospective reviews of prescribing data indicate that risperidone and olanzapine are in the top 5 prescribed medications in the U.S. and abroad, with risperidone, often the most prescribed (Bushnell et al., 2021; Klau et al., 2024; Radojcic et al., 2023).

We evaluated the long-term effects of risperidone and olanzapine in four separate experiments with female mice. We used an oral self-dosing procedure to reduce stress associated with drug administration and to ensure rapid drug consumption. After we established weight gain during treatment, we conducted extended post-treatment evaluations to give treatment-induced weight differences ample time to subside. Following the extended post-treatment evaluations, we exposed mice to a high-fat diet to evaluate susceptibility to diet-induced weight gain, long after treatment cessation. Although we evaluated the effects of risperidone in male mice (data reported in supplemental material only), all other experiments utilized female mice because the male mice failed to show treatment-induced weight gains and were therefore not useful for evaluating whether treatment-induced weight gains persisted after treatment cessation (data for male mice available in the supplemental file). The failure of male mice to exhibit SGA-induced weight gain is consistent with multiple studies in rats and mice (Albaugh et al., 2006; Choi et al., 2007; Davey et al., 2012; Pouzet et al., 2003; Seguin et al., 2023; Zapata & Osborn, 2020). Across experiments, we varied the dose of the drugs, allowing a determination of the robustness of effects. Further, across experiments, there were slight variations in the age at which treatment started and ended and the duration of the post-treatment evaluation phase and high-fat diet phase, allowing a determination of the robustness of effects against small variations in age. All data and code are available at https://osf.io/rypkq/?view_only=d2b26a237d104dbb939014bb70069703.

## Methods

### Subjects

Female (n = 84) and male (n = 16) C57BL/6J mice (#000664; Jackson Laboratory, Bar Harbor, ME) were used. Most studies were conducted using female mice based on findings in the literature that female rodents are more sensitive to gaining weight when administered SGA medications (Choi et al., 2007; Cope et al., 2005; Lord et al., 2017; Zapata & Osborn, 2020). Mice were housed individually in shoe-box style cages under a 12-h light/dark cycle with lights on at 7:00 a.m. and off at 7:00 p.m. Mice had unlimited access to standard rodent chow (Laboratory Rodent Diet 5001; LabDiet) and water in their home cages. All studies were approved by the Institutional Animal Care and Use Committee at the Pennington Biomedical Research Center and performed in accordance with the *Guide for the Care and Use of Laboratory Animals*.

### Procedure

#### Drugs

Risperidone (Item No. 13629; Cayman Chemical, Ann Arbor, MI) and olanzapine (Item No. 11937; Cayman Chemical) were administered using an oral self-dosing procedure, described below. Across experiments (see Table 1), doses of 3 mg/kg/day or 0.5 mg/kg twice daily were used for risperidone and 3 or 6 mg/kg/day olanzapine. Doses were chosen based on 1) previous studies reporting long-term changes in brain structure and function in rats exposed to 7.5 mg/kg/day olanzapine for a limited period of time early in life (Milstein et al., 2013; Vinish et al., 2013; Xu et al., 2015), 2) estimates that clinically relevant dopamine D2 receptor occupancy occurs at risperidone doses of 0.5 – 1 mg/kg/day and olanzapine doses of 1 – 2 mg/kg/day (Kapur et al., 2003), 3) higher rates of metabolism of these drugs in rodents versus humans (Kapur et al., 2003) and 4) our reasoning that because we planned to use orally administered doses, subject to first-pass metabolism, higher doses than those reported in the literature with injections might need to be given to achieve clinically relevant dopamine D2 receptor occupancy and weight gain. Importantly, we were interested in whether weight gain induced by SGA treatment persisted following the treatment period and thus our primary goal for dose selection was to use doses that produced weight gain.

**Table 1.**
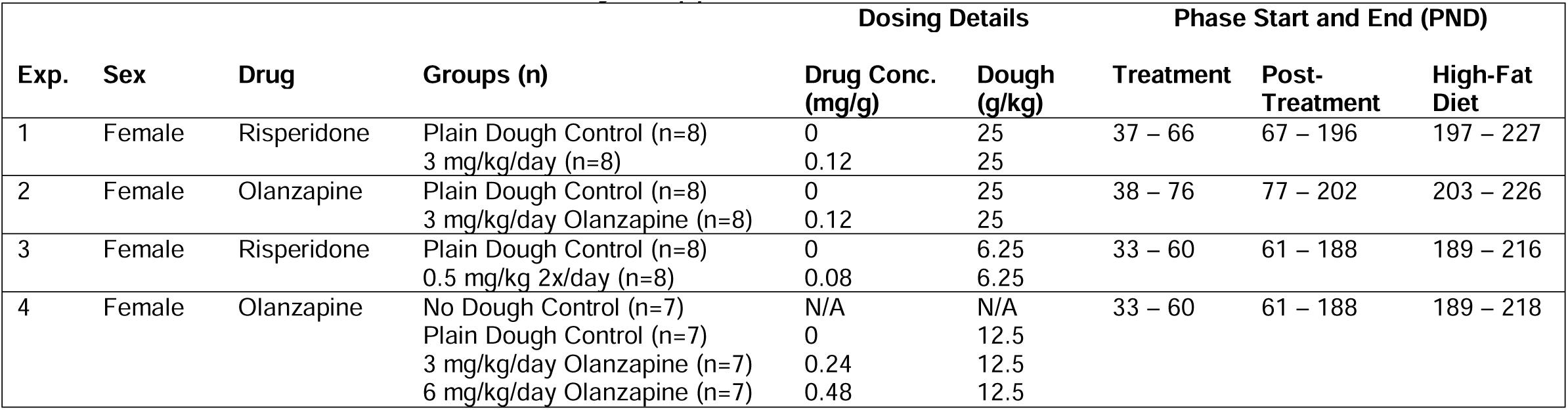
Description of experiments and ages at start and end of each phase. Details of each experiment including sex of mice, drug, groups of mice including number of mice per group, the concentration of drug in dough (mg/g), the amount of dough administered (g/kg), and the ages at which drug was administered daily (“Treatment”), the ages following the drug administration phase before mice were given access to high-fat diet (“Post-Treatment”), the ages at which mice had access to the high-fat diet, and the ages after high-fat diet was removed and mice continued in the study, if applicable.

#### Oral Self-Dosing

An oral self-dosing procedure was selected for drug administration because it provides a low stress method for chronic oral drug delivery relative to injection or oral gavage. Mice were first exposed to 3 days of dough habituation by applying ∼0.5 g of cookie dough to the inside of the rim of a 25 ml glass bottle and placing the bottle on the floor of the cage, oriented such that portion of the rim to which the dough was affixed was on the top to avoid contact with bedding. Plain cookie dough was mixed using the following proportions (based on weights) of ingredients: 1) 0.74 Betty Crocker Sugar Cookie Mix (General Mills, Inc., Minneapolis, MN), 2) 0.17 unsalted Land O’ Lakes Sweet Cream Butter (Land O’ Lakes, Inc., Arden Hills, MN), and 0.09 Eggbeaters (Con Agra Foods, Inc., Chicago, IL). By the third day of dough habituation, all mice rapidly consumed the cookie dough (< 5 min). For drug dosing, the proportion of ingredients remained the same except that the dry ingredient portion included the cookie mix and the drug (see Table 1 for details of each experiment).

#### Body Weight and Food Consumption

Mice and food in the cage hopper were weighed regularly multiple times per week for the duration of the study to assess changes in body weight and food consumption. Average daily food consumed was calculated for each measurement day as the difference between the prior and current day food weights divided by the number of intervening days. Caloric intake of diet was calculated by multiplying the caloric content of the diet (LabDiet 5001: 3.02 kcal/g; Research Diets D12451: 4.73 kcal/g), provided on the product information sheet, by the grams of diet consumed. Caloric intake of cookie dough was calculated by multiplying the estimated kilocalories per gram of dough (4.17 kcal/g) by the grams of dough consumed. The kilocalories per gram of dough was estimated as the sum of the products of the kilocalories per gram provided by each ingredient and the proportion of each ingredient in dough (Butter: 7.14 kcal/g x 0.17 = 1.21 kcal/g of dough; Egg Beaters: 0.54 kcal/g x 0.09 = 0.05 kcal/g of dough; Sugar Cookie Mix: 3.93 kcal/g x 0.74 = 2.91 kcal/g of dough), based on the information on each ingredient product label.

#### High-Fat Diet

The effects of consuming a high-fat diet providing 45% kcal fat, 35% kcal carbohydrate, and 20% kcal protein (D12451; Research Diets, Inc., New Brunswick, NJ) on body weight were evaluated following the post-treatment phase (see Table 1). Mice and food were weighed daily to assess changes in body weight and high-fat diet consumption. Daily consumption of high-fat diet for each measurement day was calculated as the difference between two consecutive food weights divided by the number of days between the two consecutive measurements (except in rare cases, mice and food were weighed every day). Caloric intake of the high-fat diet was calculated by multiplying the grams of diet consumed by the caloric intake of the diet (4.73 kcal/g).

#### Tissue and Blood Collection

After a 4 hour fast, mice from Experiments 3 (0.5 mg/kg twice daily risperidone-treated and vehicle control female mice) and 4 (3 and 6 mg/kg/day olanzapine-treated, vehicle control, and no dough female mice) were euthanized via CO2 asphyxiation and then decapitated (no tissues or blood was collected from mice in Experiments 1 and 2). Immediately following decapitation, blood samples were collected via trunk blood and the serum fraction was separated for downstream analysis. Gonadal white adipose tissue (gWAT) samples were removed and frozen in liquid nitrogen.

#### RNA Expression, Serum Insulin, and Blood Glucose

RNA from adipose tissue and gene expression was assessed as described previously (Collier et al., 2021). Briefly, RNA was isolated from adipose tissue using the RNeasy Mini kit (Qiagen) per the manufacturer’s instructions. cDNA synthesis was conducted using the iScript kit (Bio-Rad). Gene expression was detected using cDNA as a template in a SYBR Green real-time PCR reaction on a CFX96 Touch Real-Time PCR Detection System (Bio-Rad; primer sequences are provided in Table S1). Genes tested by RT-PCR were normalized using ribosomal protein S9 (*Rps9*) as a housekeeping gene. Each sample was run in duplicate and analyzed by the delta-delta Ct method. Serum insulin was measured using the Mouse Insulin ELISA kit from Mercodia (Uppsala, Sweden) according to the manufacturer’s instructions. Blood glucose was measured using a Bayer Contour Glucometer.

### Data Analysis

Changes in body weight and kilocalories (kcal) consumed were analyzed using a multilevel linear model analysis, which is a linear regression approach that estimates group- and individual-level coefficient estimates. Multilevel models are increasingly recognized as providing benefits over repeated measures analysis of variance (ANOVA) when analyzing data for which multiple observations have been collected within individual animals, when the same animals are used across conditions, and when observations are unbalanced (Boisgontier & Cheval, 2016; de Melo et al., 2022; Yu et al., 2022) Further, a linear regression approach increases power when using continuous predictors such as time because linear regression approaches do not categorize continuous variables like ANOVA does and therefore utilize fewer degrees of freedom (Lazic, 2008; Lazic, 2018). Multilevel linear models were conducted using the lmerTest pack in R (Kuznetsova et al., 2017), which provides p-values for linear mixed effects models created by the lmer function of the lme4 package (Bates et al., 2015).

Within each experiment, body weight was analyzed using fixed effect predictors of age (PND; continuous and transformed so that the first day of each phase was set to 0), Group (categorical, dummy-coded with plain dough group as the reference level), Phase (categorical, dummy coded with treatment phase as the reference level; treatment, post-treatment, and high-fat diet) and all 2- and 3-way interactions of those predictors. The random effects portion of each model was defined using a “complex random intercept” approach with an estimate for the coefficient of age specified for the combination of Mouse and Phase and Mouse alone, based on a recent paper suggesting the use of complex random intercepts to reduce Type I error inflation when conducting multilevel model analyses of fully crossed designs (Scandola & Tidoni, 2024). Caloric intake was analyzed in the same way as body weight. The emmeans and emtrends commands of the emmeans package (Lenth, 2020) were then used to calculate fixed-effect level estimates for the intercept and coefficient of PND, respectively, for each Group within each Phase and to assess differences between Groups for statistical significance. A separate model was conducted for each combination of outcome and experiment. Finally, to estimate the overall effect of early-life SGA treatment on body weight and caloric intake in female mice, a combined analysis was conducted for each outcome (body weight and caloric intake) in which the data from all 4 experiments were analyzed together. For the combined analysis, all treated mice were labeled as “Treatment,” all plain dough control mice as “Control,” and the no dough group from Experiment 4 was not included in the analysis. Otherwise, the combined analysis was conducted as described for the individual experiment analyses.

For statistical analysis of gene expression data collected in Experiments 3 (0.5 mg/kg risperidone twice daily) and 4 (3 and 6 mg/kg/day olanzapine), 2^—ΔΔCT^ values were log-transformed and analyzed using a general linear model with the lm() command in R. A separate analysis was conducted for each experiment and category of genes (Inflammation-associated genes: *Adgre*, *Ccl2*, *Cd68*, *Itgam*; adipogenesis-associated genes: *Fabp4*, *Leptin*; and lipogenesis-associated genes: *Acaca*, *Fasn*) using gene (within the category; dummy-coded with the first gene in each category set as the reference) and group (dummy-coded with the plain dough control group as the reference) as factors. Following each analysis, post-hoc pairwise comparisons between groups in the experiment were conducted using the emmeans command of the emmeans package.

Serum insulin concentration (ng/mL) and blood glucose concentration (mg/dL) from mice used in Experiments 3 and 4 were analyzed using a general linear model via the lm() command in the R programming language with group (dummy-coded with plain dough control as reference) as a predictor. A separate analysis was conducted for each experiment. Post-hoc pairwise comparisons between groups in the experiment were conducted using the emmeans command.

Data and model visualizations were conducted using the ggplot2 package (Wickham, 2016) in the R programming language.

## Results

### Effects of Early Adolescent SGA Administration on Body Weight and Caloric Intake

At the beginning of the treatment phase, the intercepts (estimated starting body weight) for all groups of mice averaged 15.6 g (range 15.2 – 16.1 g; Figure 1A-D; for complete model results see supplemental file). Differences in estimated starting body weights between treated and control groups averaged 0.06 g (range: -0.39 to 0.73, p-value range: 0.202 – 0.997; Figure 1E, Tx panel). During the treatment phase, the average rate of weight gain across all groups was 0.10 g/day (range: 0.05 – 0.14 g/day; Figure 1A-D), with differences in estimated rates of weight gain between female mice treated with risperidone or olanzapine and their respective controls averaging 0.04 g/day (range 0.03 – 0.05 g/day, p-value range: 0.017 – 0.211; Figure 1F, Tx panel). The estimated weight for the no dough group at the start of the treatment phase was the same as that of the control group (15.6 g in both groups; Figure 1D) and the no dough group gained weight at a rate higher than that of the plain dough control group (0.10 vs. 0.08 g/day; p = 0.708; Figure 1F, Tx panel).

**Figure 1.**
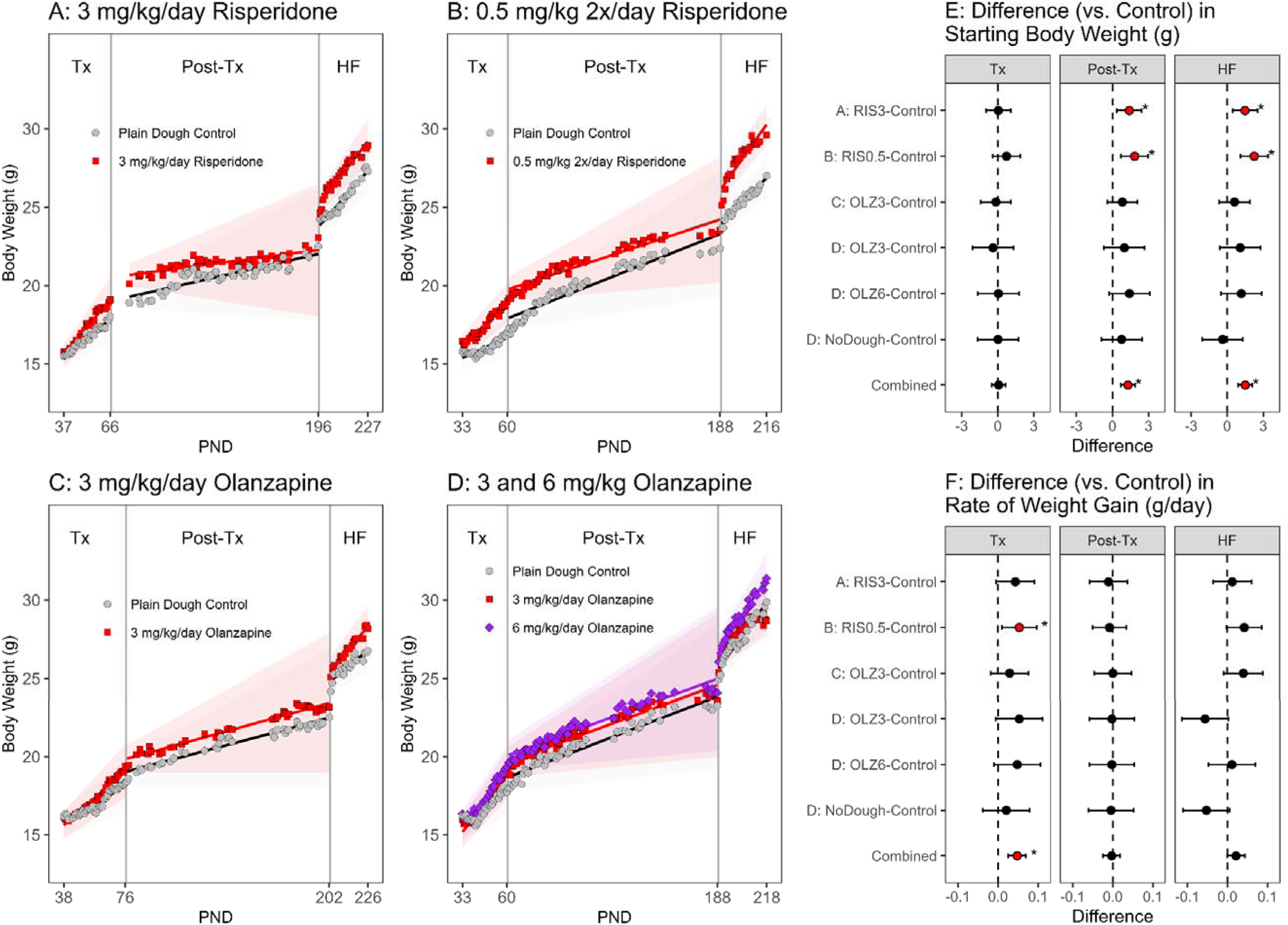
Daily body weight. Average body weight (g) across days during the treatment (Tx), post-treatment (Post-Tx), and high-fat diet (HF) phases. **A**: Results from the mice (n = 8/group) fed either plain cookie dough daily or dough with risperidone daily for a target dose of 3 mg/kg/day. **B**: Results from mice (n = 8/group) fed either plain cookie dough twice daily or dough with risperidone twice daily for a target dose of 0.5 mg/kg twice daily. **C**: Results from mice (n = 8/group) fed either plain cookie dough daily or dough with olanzapine daily for a target dose of 3 mg/kg/day. **D**: Results from mice (n = 7/group) fed either plain cookie dough daily, dough with olanzapine daily for a target dose of 3 mg/kg/day, and dough with olanzapine daily for a target dose of 6 mg/kg/day (results from a group of mice not fed any dough are not plotted for sake of clarity). **A-D**: Symbols represent group average values. Straight lines indicate regression model predictions and shaded regions around the straight lines indicate the 95% confidence limits of the predictions. **E**: Differences between the model-estimated intercept values for each group and the corresponding value for the plain dough control mice. Error bars represent 95% confidence limits. Asterisks indicate differences with p < 0.05. The “Combined” estimates are for model-estimated parameters obtained from a model fitted to the data from all 4 experiments. **F**: Differences between model-estimated slope values in each group and the corresponding value for the plain dough control mice. Other details as in E.

At the beginning of the treatment phase, female mice consumed an average of 12.2 kcal per day (combined food and cookie dough; range: 11.5 – 13.3), as estimated by averaging the model intercepts for all groups (Figure 2A-D). Differences between the treated groups and their respective control groups in starting caloric intake were small relative to overall intake (average difference = 0.66 kcal; range: 0.14 – 1.49; p-value range: 0.185 – 0.876; Figure 2E, Tx panel). Across days in the treatment phase, caloric intake declined by an average of 0.03 kcal/day (range: -0.13 to 0.09 kcal/day), with differences between treated groups and respective control groups on average equal to 0.03 kcal/day (range: -0.02 to 0.07 kcal/day, p-value range: 0.102 – 0.874; Figure 2F, Tx panel). Mice in the no dough group consumed an estimated 12.6 kcal/day at the start of the treatment phase (Figure 2D), which was 0.52 kcal greater than the estimated starting consumption value in the control group (Figure 2E, Tx panel; p = 0.880). Across days in the treatment phase, intake increased in the no dough group (0.09 kcal/day) whereas caloric intake decreased in the plain dough control group (-0.02 kcal/day; difference between groups = 0.12 kcal/day, p = 0.035; Figure 2F, Tx panel).

**Figure 2.**
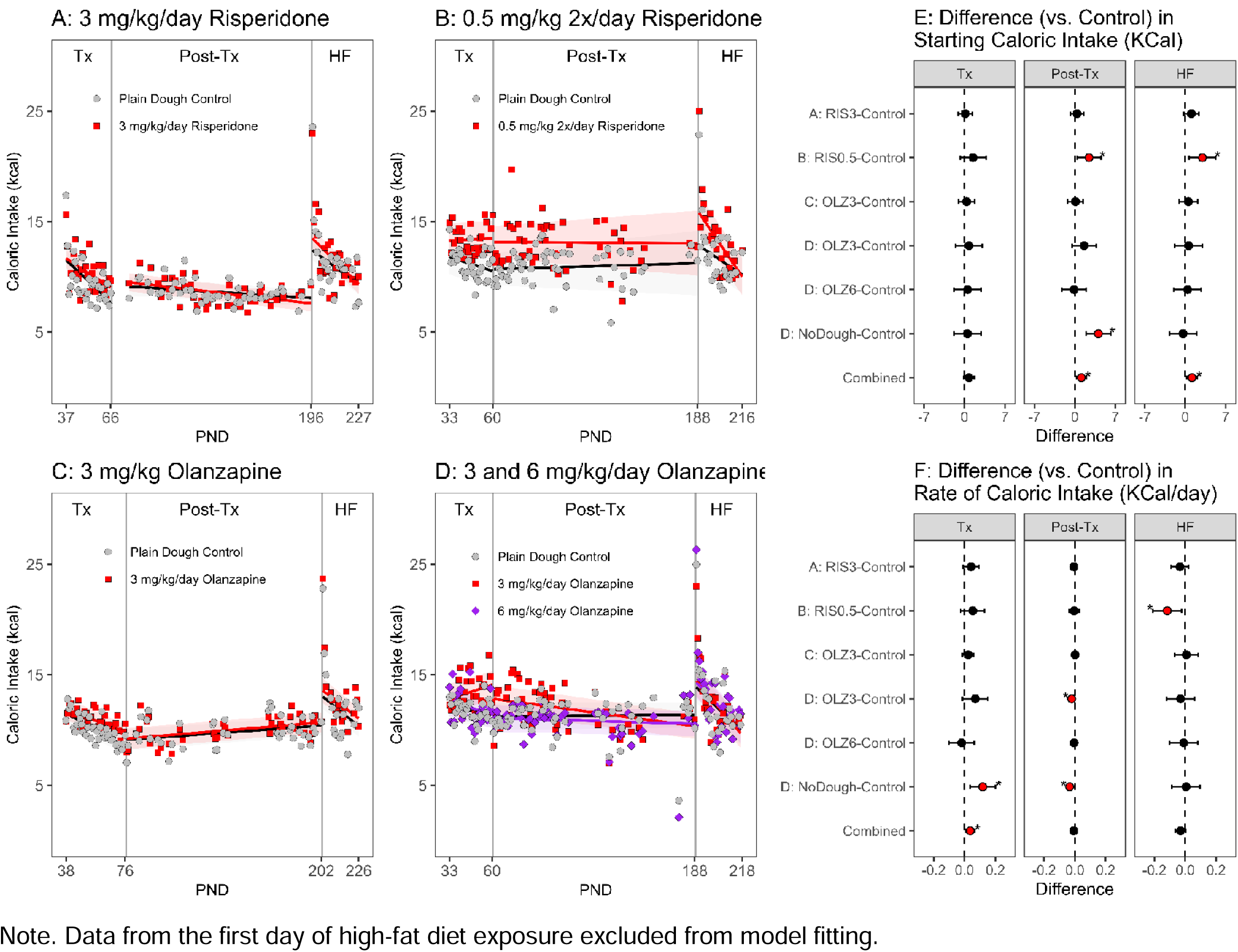
Daily caloric intake. Average caloric intake in kilocalories across days during the treatment (Tx), post-treatment (Post-Tx), and high-fat diet (HF) phases. **A**: Results from the mice (n = 8/group) fed either plain cookie dough daily or dough with risperidone daily for a target dose of 3 mg/kg/day. **B**: Results from mice (n = 8/group) fed either plain cookie dough twice daily or dough with risperidone twice daily for a target dose of 0.5 mg/kg twice daily. **C**: Results from mice (n = 8/group) fed either plain cookie dough daily or dough with olanzapine daily for a target dose of 3 mg/kg/day. **D**: Results from mice (n = 7/group) fed either plain cookie dough daily, dough with olanzapine daily for a target dose of 3 mg/kg/day, or dough with olanzapine daily for a target dose of 6 mg/kg/day (results from a group of mice not fed any dough are not plotted for sake of clarity). **A-D**: Symbols represent group average values. Straight lines indicate model predictions and shaded regions around the straight lines indicate the 95% confidence limits of the predictions. **E**: Differences between the model-estimated intercept values for each group and the corresponding value for the plain dough control mice. Error bars represent 95% confidence limits. Asterisks indicate differences with p < 0.05. The “Combined” estimates are for model-estimated parameters obtained from a model fitted to the data from all 4 experiments. **F**: Differences between model-estimated slope values in each group and the corresponding value for the plain dough control mice. Other details as in E.

### Post-Treatment Effects of Early Adolescent SGA Administration on Body Weight and Caloric Intake

At the beginning of the post-treatment phase, the average estimated starting weight was 19.5 g (range: 17.9 – 20.7 g; Figure 1A-D). Estimates of starting weight were consistently higher in previously treated female mice than their respective controls, with differences between treated and control groups across experiments averaging 1.29 g (range: 0.83 – 1.84 g, p-value range: 0.002 – 0.362; Figure 1E, Post-Tx panel).

Across days in the post-treatment phase, body weights increased, on average, 0.03 g/day (range: 0.01 – 0.04 g/day; Figure 1A-D) with small differences in slopes between treated and control groups (average difference = -0.004; range: -0.010 – 0.001, p-value range: 0.675 – 0.997; Figure 1F, Post-Tx panel). Mice in the no dough group started the post-treatment phase at an estimated 19.5 g, slightly heavier than the control group (difference = 0.76 g; p = 0.555) and gained weight during the post-treatment phase at a lower rate than the control group (difference = -0.004 g/day, p = 0.986; Figure 1F, Post-Tx panel).

At the beginning of the post-treatment phase, mice consumed, on average, 11.1 kcal per day (range: 9.1 – 15.3 kcal/day; Figure 2A-D), with differences between treated and control groups ranging averaging 0.88 kcal/day (range: -0.14 – 2.45 kcal/day; p-value range: 0.022 – 0.989; Figure 2E, Post-Tx panel). Over the duration of the post-treatment phase, caloric intake declined gradually in some groups and increased gradually in others (average = -0.01, range: -0.03 – 0.01 kcal/day; Figure 2A-D), with small differences between treated and control groups averaging -0.01 kcal/day (range: - 0.02 – 0.001 kcal/day, p-value range: 0.038 – 0.871; Figure 2F, Post-Tx panel). In comparison, caloric intake declined over the post-treatment phase in the no dough control group and increased in the plain dough control group (-0.03 vs. 0.00 kcal/day, p < .001; Figure 2F, Post-Tx panel).

### High-Fat Diet-Induced Weight Gain in Adulthood

At the beginning of the high-fat diet phase, the average starting body weight was 25.4 g (range: 23.9 – 26.8; Figure 1A-D), with an average difference between treated and control groups of 1.32 g (range: 0.61 to 2.25 g, p-value range: 0.0003 – 0.328; Figure 1E, HF panel). In comparison, the average starting body weight in the no dough group was 25.3 g, 0.38 g less than the average body weight in the control group (Figure 1F, HF panel). During the high-fat diet phase, mice gained, on average, 0.12 g/day (range: 0.08 – 0.16 g/day; Figure 1A-D). Differences between treated and control groups in rate of weight gain averaged 0.01 g/day (range: -0.06 – 0.04, p-value range: 0.058 – 0.914; Figure 1F, HF panel). In comparison, the no dough control group gained weight at a rate of about 0.05 g/day less than the plain dough control group (Figure 1F, HF panel).

### Combined Model Analysis of Body Weight and Caloric Intake

In the combined model analysis of body weight across experiments, body weight of treated mice was not greater at the start of the Treatment Phase, but was greater at the start of the Post-Treatment and High-Fat Diet Phases with an estimated difference of 1.29 and 1.50 grams, respectively (Figure 1E, Tx Panel). During the Treatment Phase, the estimated rate of weight gain was higher in treated mice during the treatment phase (0.12 vs. 0.07 g/day, p < 0.001; Figure 1F, Tx Panel), equal in the Post-Treatment Phase (0.03 g/day, p=0.798; Figure 1F, Tx Panel), and slightly higher during the High-Fat Diet Phase (0.13 vs. 0.11 g/day, p = 0.062; Figure 1F, Tx Panel).

In the combined model analysis of caloric intake across experiments, starting caloric intake differences between treated and control mice increased across phases, starting at a difference of 0.76 kcal, then 1.12 kcal, and finally 1.28 kcal during the Treatment, Post-Treatment, and High-Fat Diet Phases (p = 0.098, 0.010, and 0.008, respectively; Figure 2E, Tx Panel). The rate of change in caloric intake across days was negative for both treated and control mice in the Treatment phase but was declined less per day in treated mice (-0.04 vs. -0.08 kcal/day, p = 0.005; Figure 2F, Tx Panel). The rate of change in caloric intake across days in the Post-Treatment phase was 0 in both treated and control mice (Figure 2F, Tx Panel), and was negative in both treated and control groups during the High-Fat Diet Phase (-0.14 vs. -0.10 kcal/day, p = 0.029; Figure 1F, Tx Panel).

### Circulating Insulin and Glucose

Serum insulin concentration at the end of the study was not altered by prior early adolescent risperidone (Figure S2A) or olanzapine (Figure S2B) treatment. Similarly, blood glucose concentration at the end of the study was not altered by prior early adolescent risperidone (Figure S2C) or olanzapine (Figure S2D) treatment.

### Gene Expression in gWAT

Gene expression in gonadal white adipose tissue (gWAT) was measured in the mice from Experiments 3 and 4 to assess whether early adolescent SGA treatment produces long-term changes in expression of inflammation-, adipogenesis-, and lipogenesis-associated genes. In risperidone-treated and control female mice, relative expression levels of inflammation-, adipogenesis-, and lipogenesis-associated genes were equivalent between treatment and control groups (Figure 3A-C). In contrast, in olanzapine-treated and control female mice, the 6 mg/kg/day group exhibited greater expression in 3 of 4 inflammation-associated genes, 1 of 2 adipogenesis-associated genes, and 2 of 2 lipogenesis-associated genes (Figure 3D-F). The 3 mg/kg/day group exhibited an increased expression in 2 of 2 lipogenesis-associated genes (Figure 3F).

**Fig 3.**
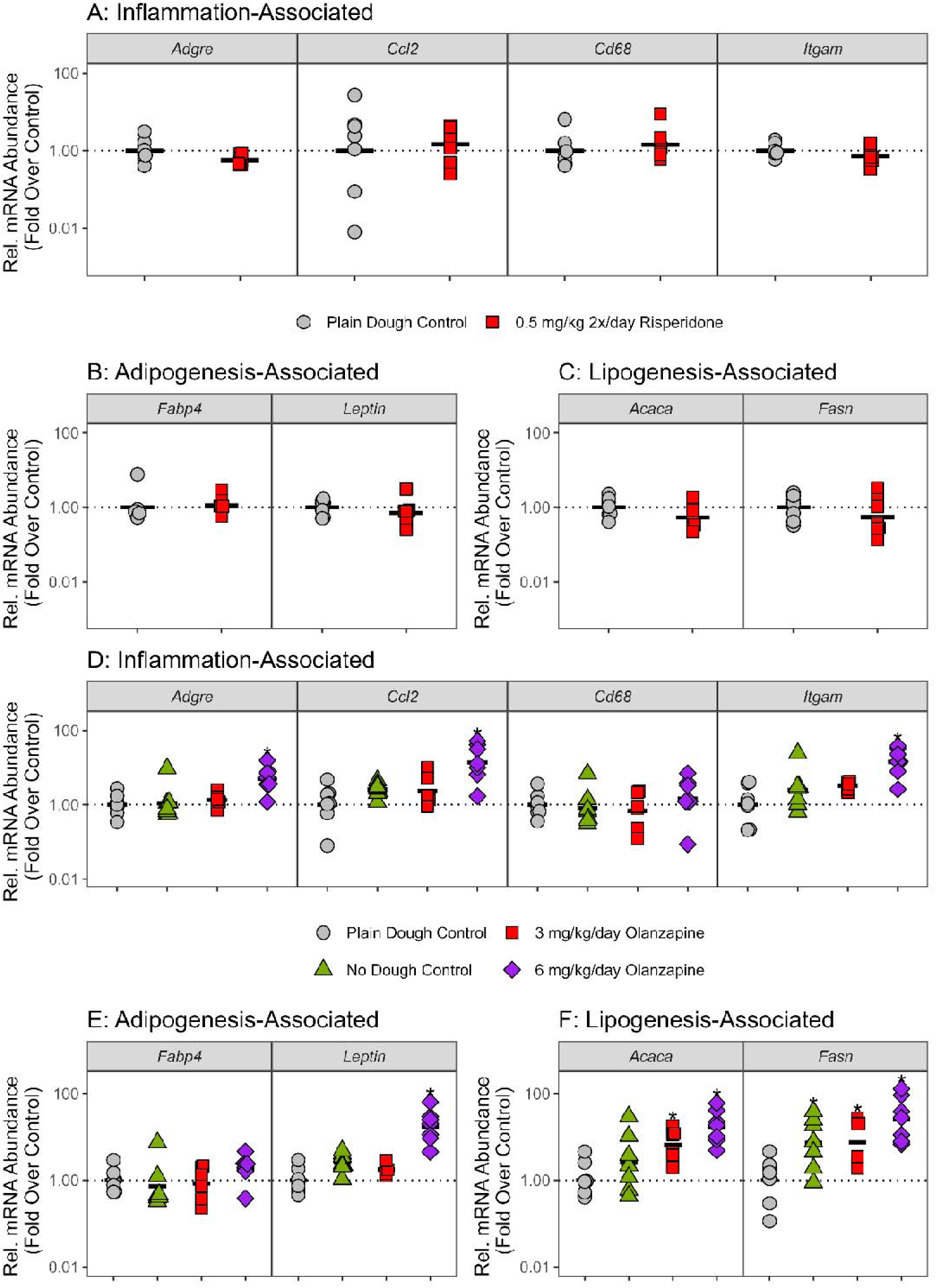
gWAT mRNA abundance in female mice. Relative mRNA abundance levels in gonadal white adipose tissue (gWAT) calculated using the delta delta cycle threshold method for female mice previously treated with risperidone or vehicle for **A**: inflammation-associated genes; **B**: adipogenesis-associated genes; and **C**: lipogenesis-associated genes and for female mice previously treated with olanzapine or plain dough or who were not given any cookie dough (“No Dough Control”) for **D**: inflammation-associated genes; **E**: adipogenesis-associated genes; and **F**: lipogenesis-associated genes. Black horizontal bars represent group means. Each individual data point represents the result obtained in an individual mouse. Asterisks indicate differences with p < 0.05

## Discussion

The current study examined the acute and post-treatment effects of early adolescent exposure to risperidone and olanzapine on body weight and caloric intake. At the end of the study, some mice were used to assess circulating glucose and insulin and expression of genes associated with inflammation, adipogenesis, and lipogenesis. Across experiments, early adolescent treatment with risperidone or olanzapine produced weight gains that resulting in body weights at the start of the Post-Treatment Phase between 4.4 and 10.3% above controls and those weight gains persisted for months following treatment cessation with body weights of treated mice between 2.4 and 9.4% above controls at the start of the High-Fat Diet Phase. The results from individual experiments indicating persistent weight gains induced by adolescent treatment with risperidone and olanzapine were supported by a combined analysis using all the data showing treated mice had body weights of 6.9 and 6.1% at the start of the Post-Treatment and High-Fat Diet Phases. There was little evidence to support a conclusion of increased susceptibility to diet-induced weight gain with rates of weight gain during high-fat diet feeding only marginally higher in treated mice compared to controls in some experiments and this was consistent with the combined analysis showing that the overall difference in rate of weight gain between treated and control mice was 0.02 g/day (p = 0.062). In Experiment 4, mice previously treated with olanzapine exhibited increased expression of inflammatory-, adipogenesis-, and lipogenesis-associated genes in adulthood, long after treatment cessation. Together, these results strongly suggest that body weight increases and gene expression changes induced by early adolescent SGA treatment can persist long after treatment cessation.

The pharmacological target of SGA medications that underlies treatment-induced weight gain remains unclear but may involve multiple targets of these medications and the actions of these medications may interact with other variables such as diet and genetic factors (see De Hert, Detraux, et al., 2011 for a discussion). Multiple neurotransmitter systems, including serotonin, histamine, and dopamine have been implicated in the weight increasing effects of SGA medications (De Hert, Detraux, et al., 2011). For example, pharmacological blockade or deletion of 5-HT_2C_ receptors has been reported to ameliorate the weight gain inducing effects of olanzapine in mice (Lord et al., 2017). Similarly, genetic deletion of melanocortin 4 receptors in Sim1-expressing neurons has been reported to prevent risperidone-induced weight gain and treatment with a melanocortin 4 receptor agonist has been reported to prevent risperidone- and olanzapine-induced weight gain (Li et al., 2021). Thus, it appears likely that multiple receptor targets may confer risk of weight gain.

In the current study, persistent increases in body weight do not appear to be driven solely by differences in caloric intake because differences in caloric intake were small and further, in Experiment 4, the No Dough group exhibited increased caloric intake but failed to exhibit the increased body weight seen in treated mice. That SGA treatment was associated with increased weight gain without increased caloric intake is consistent with other studies (Bahr et al., 2015; Zapata & Osborn, 2020). Yet, other studies have reported that SGA treatment was associated with increased food intake and in some cases that increased food intake was necessary for the increased body weight gain (Li et al., 2021; Li et al., 2013; Lord et al., 2017). Because SGA treatment can suppress activity levels and energy expenditure (Bahr et al., 2015; Cope et al., 2009; Lord et al., 2017), body weight can increase even if food intake does not, but the factors that determine when food intake increases are necessary for effects on body weight remain unclear.

We expected that mice treated with risperidone or olanzapine would exhibit increased weight gain on a high-fat diet based on findings that treatment with risperidone or olanzapine can induce increases in expression of dopamine D2 receptors that persist following treatment cessation (De Santis et al., 2016b; Milstein et al., 2013; Vinish et al., 2013) and the finding that early adolescent overexpression of dopamine D2 receptors resulted in a subsequent increased sensitivity to high-fat diet-induced weight gain in adulthood despite prior normalization of D2 receptor expression (Labouesse et al., 2018). Evidence for a persistent effect of previous SGA treatment on diet-induced sensitivity was weak in the current study. At the individual experiment level, 2 experiments (0.5 mg/kg risperidone twice daily and 3 mg/kg/day olanzapine) exhibited rates of body weight increase that were positive with 95% confidence limits that included 0. Similarly, in the combined analysis, the difference in rate of weight gain between treated and control mice was 0.02 g/day with 95% confidence limits that just barely included 0 (-0.001 – 0.043). Thus, the current results present only weak evidence for the possibility that SGA treatment may produce increased sensitivity to diet-induced weight gain.

The finding that early adolescent exposure to second-generation antipsychotic medications can produce long-term effects on body weight that persist well beyond the treatment period is consistent with a number of studies in rodents reporting long-term changes in brain structure and function, behavior, and responses to drugs, following acute exposure to SGA medications early in life (Bardgett et al., 2019; Bardgett et al., 2020; Bardgett et al., 2013; Bardgett et al., 2024; De Santis et al., 2018; De Santis et al., 2016a, 2016b; Frost et al., 2010; Gannon et al., 2015; Kendricks et al., 2024; Lee Stubbeman et al., 2017; Milstein et al., 2013; Vinish et al., 2013; Xu et al., 2015).

Interestingly, long-term changes in some of these studies often occurred in the absence of body weight increases induced by treatment (e.g., Bardgett et al., 2013; Kendricks et al., 2024; Vinish et al., 2013). Similarly, we previously reported long-term changes in body composition and metabolic outcomes in male and female mice fed a high-fat diet and treated with olanzapine, despite an absence of treatment-induced increases in weight gain (Soto et al., 2022). That long-term changes in some outcomes can occur without treatment-induced increases in weight gain suggests that the mechanisms underlying different long-term outcomes may be distinct.

In the current study, our primary purpose was to evaluate how long weight gain induced by SGA treatment persists following treatment cessation. Our purpose was informed by the well-established findings that many, if not most, SGA medications induce weight gain in children and adults (Dayabandara et al., 2017; van der Esch et al., 2021). Thus, our focus was on the selection of doses of risperidone and olanzapine that would reliably induce weight gain. Importantly, the weight gain we observed during treatment (4.4 – 10.3% of controls) reached or exceeded clinically significant levels of 7% (De Hert, Detraux, et al., 2011) in 3 of 5 comparisons of treated and control female mice (treated vs. control in Experiments 1 and 3 and 6 mg/kg olanzapine vs. control in Experiment 4). It is unclear to what degree the doses used are equivalent to doses used in pediatric patients. Although a more precise delineation of the equivalence of doses in mice and humans would be useful, the matter is complicated by questions regarding the basis on which to equate doses (cf. Kapur et al., 2003). Still, the present results are relevant to the question of SGA-induced long-term weight gain because a key aspect of the issue is what happens once SGA-induced weight gain occurs. To that extent, the current results demonstrate that weight gain produced during treatment remains indefinitely or dissipates very slowly following treatment cessation.

The multilevel modeling approach used in the current study may be unfamiliar to some readers and therefore warrants some discussion. Although more complex than a repeated measures analysis of variance (RMANOVA), the multilevel modeling approach is better suited for longitudinal, repeated measures data and offers several advantages (Boisgontier & Cheval, 2016; de Melo et al., 2022; Ma et al., 2012; Yu et al., 2022).

RMANOVA incurs inflated Type I error rates when conducted on data that are nested (multiple observations per animal) and crossed (observations collected from the same animal in each condition), whereas multilevel models account for nested and crossed nature of data and provide acceptable Type I error rates (Boisgontier & Cheval, 2016). Further, RMANOVA also does not deal well with missing data - in a RMANOVA, missing observations result in dropping of all data for the affected case (e.g., missing body weights for a mouse on some occasions cause the removal of all data for the affected mouse) whereas multilevel models are robust against missing/unbalanced data (Boisgontier & Cheval, 2016). Finally, regular ANOVA and RMANOVA treats continuous variables as categorical leading to multiple disadvantages such as the need for multiple post-hoc comparisons at each value of the predictor variable (e.g., compare groups at time point 1, at time point 2, etc.; within a group, compare outcome at time points 1 and 2, at time points 2 and 3, etc.) and the opportunity for any deviation from equality, even when not systematic, to drive statistical significance and interpretative resources (Young, 2016). Treating continuous variables as continuous simplifies analysis and interpretation and promotes parsimony (Lazic, 2008; Young, 2016).

In conclusion, the current results demonstrate that early adolescent exposure to SGA medications can produce long-lasting increases in body weight and expression of inflammation-, adiposity-, and lipogenesis-associated genes. Further, these results suggest that early adolescent SGA exposure confers a long-term increased sensitivity to diet-induced weight gain. The mechanism underlying these long-term changes may involve overexpression of dopamine D2 receptors although further research is needed to address this question. An understanding of the underlying mechanism(s) will be important for the design of medications that provide therapeutic value with reduced long-term undesirable consequences.

## Notes

https://osf.io/rypkq/?view_only=d2b26a237d104dbb939014bb70069703

### Competing Interest Statement

The authors have declared no competing interest.

