## Supplemental Tables and Figures for "Long-Term Effects of Early Adolescent Second-Generation Antipsychotic Exposure on Body Weight, Caloric Intake, and Metabolism-Associated Gene Expression"

### Supplemental Material

**Supplemental Tables**

Table S1. Primer sequences.
Table S2. Regression coefficients from body weight models
Table S3. Regression coefficients from caloric intake models
Table S4. Intercepts and slopes for body weight
Table S5. Intercepts and slopes for caloric intake

**Supplemental Figures**

Figure S1. Body weight and caloric intake in male mice
Figure S2. Circulating insulin and glucose in female mice

Table S1. Primer sequences for RT-PCR.

| Gene | F Primer (5' - 3') | R Primer (5' - 3') |
|---|---|---|
| *Adgre* | TGCATCTAGCAATGGACAGC | GCCTTCTGGATCCATTTGAA |
| *Ccl2* | CCCACTCACCTGCTGCTACT | TCTGGACCCATTCCTTCTTG |
| *Cd68* | TCCAAGCCCAAATTCAAATC | ATTGTATTCCACCGCCATGT |
| *Itgam* | GACTCAGTGAGCCCCATCAT | AGATCGTCTTGGCAGATGCT |
| *Fabp4* | CATCAGCGTAAATGGGGATT | TCGACTTTCCATCCCACTTC |
| *Leptin* | TGACACCAAAACCCTCATCA | TCATTGGCTATCTGCAGCAC |
| *Acaca* | GGTCTTCGAGTGGATTGGCA | ATCCCTTTCCCTCCTCCTCC |
| *Fasn* | CAAGTGTCCACCAACAAGCG | GGAGCGCAGGATAGACTCAC |

Table S2. Regression estimates from models of body weight.

Model term number, predictor, coefficient estimates (Estimate) of the predictor, standard errors (SE) of the estimates, and resulting t-statistic (t), degrees of freedom (df; Satterthwaite estimated), and p-values (p) from the multilevel linear regression analysis of body weight. The fixed effects portion of the model included estimates for the predictors Group (dummy-coded with the plain dough control group as the reference level for all cohorts), Phase (dummy-coded with the Treatment phase as the reference level), and PND (continuous and centered), and all possible interactions of the four predictors. The random effects portion of the model allowed the coefficient of Phase to vary by mouse.

| Model Term | Predictor | Estimate | SE | t | df | p |
|---|---|---|---|---|---|---|
| | EXP01: 3 mg/kg Risperidone | | | | | |
| 1 | (Intercept) | 18.183 | 0.762 | 23.85 | 29.25 | < .001 |
| 2 | PND | 0.081 | 0.017 | 4.87 | 38.09 | < .001 |
| 3 | Group [Treated] | 1.518 | 1.078 | 1.41 | 29.25 | .170 |
| 4 | Phase [Post-Tx] | 1.877 | 0.785 | 2.39 | 30.58 | .023 |
| 5 | Phase [HF] | 9.539 | 0.789 | 12.09 | 31.20 | < .001 |
| 6 | PND*Group [Treated] | 0.044 | 0.024 | 1.86 | 38.09 | .070 |
| 7 | PND*Phase [Post-Tx] | -0.058 | 0.021 | -2.82 | 35.38 | .008 |
| 8 | PND*Phase [HF] | 0.035 | 0.021 | 1.65 | 37.92 | .107 |
| 9 | Group [Treated]*Phase [Post-Tx] | -0.445 | 1.110 | -0.40 | 30.58 | .691 |
| 10 | Group [Treated]*Phase [HF] | 0.386 | 1.116 | 0.35 | 31.20 | .732 |
| 11 | PND*Group [Treated]*Phase [Post-Tx] | -0.054 | 0.029 | -1.85 | 35.38 | .073 |
| 12 | PND*Group [Treated]*Phase [HF] | -0.031 | 0.030 | -1.06 | 37.92 | .296 |
| | EXP03: 0.5 mg/kg RIS 2x/daily | | | | | |
| 1 | (Intercept) | 16.752 | 0.638 | 26.24 | 34.60 | < .001 |
| 2 | PND | 0.049 | 0.015 | 3.18 | 39.58 | .003 |
| 3 | Group [Treated] | 2.248 | 0.903 | 2.49 | 34.61 | .018 |
| 4 | Phase [Post-Tx] | 2.384 | 0.741 | 3.22 | 29.38 | .003 |
| 5 | Phase [HF] | 10.260 | 0.747 | 13.74 | 30.22 | < .001 |
| 6 | PND*Group [Treated] | 0.054 | 0.022 | 2.49 | 39.60 | .017 |
| 7 | PND*Phase [Post-Tx] | -0.006 | 0.020 | -0.32 | 33.75 | .753 |
| 8 | PND*Phase [HF] | 0.059 | 0.021 | 2.83 | 38.46 | .007 |
| 9 | Group [Treated]*Phase [Post-Tx] | -0.611 | 1.048 | -0.58 | 29.38 | .564 |
| 10 | Group [Treated]*Phase [HF] | 1.200 | 1.056 | 1.14 | 30.22 | .265 |
| 11 | PND*Group [Treated]*Phase [Post-Tx] | -0.061 | 0.028 | -2.15 | 33.77 | .038 |
| 12 | PND*Group [Treated]*Phase [HF] | -0.011 | 0.029 | -0.38 | 38.47 | .704 |
| | EXP02: 3 mg/kg Olanzapine | | | | | |
| 1 | (Intercept) | 18.413 | 0.915 | 20.12 | 34.18 | < .001 |
| 2 | PND | 0.063 | 0.017 | 3.79 | 35.79 | < .001 |
| 3 | Group [Treated] | 1.103 | 1.294 | 0.85 | 34.18 | .400 |
| 4 | Phase [Post-Tx] | 1.797 | 1.086 | 1.65 | 32.40 | .108 |
| 5 | Phase [HF] | 10.027 | 1.100 | 9.12 | 34.04 | < .001 |
| 6 | PND*Group [Treated] | 0.030 | 0.023 | 1.27 | 35.79 | .212 |
| 7 | PND*Phase [Post-Tx] | -0.035 | 0.023 | -1.56 | 33.95 | .128 |
| 8 | PND*Phase [HF] | 0.022 | 0.023 | 0.94 | 38.00 | .352 |

| | | | | | | |
|---|---|---|---|---|---|---|
| 9 | Group [Treated]*Phase [Post-Tx] | -0.232 | 1.536 | -0.15 | 32.40 | .881 |
| 10 | Group [Treated]*Phase [HF] | 1.211 | 1.555 | 0.78 | 34.04 | .442 |
| 11 | PND*Group [Treated]*Phase [Post-Tx] | -0.029 | 0.032 | -0.90 | 33.95 | .375 |
| 12 | PND*Group [Treated]*Phase [HF] | 0.010 | 0.033 | 0.32 | 38.00 | .750 |
| | EXP04: 3 and 6 mg/kg Olanzapine | | | | | |
| 1 | (Intercept) | 18.032 | 0.811 | 22.22 | 58.67 | < .001 |
| 2 | PND | 0.082 | 0.017 | 4.81 | 69.96 | < .001 |
| 3 | Group [No Dough] | 0.641 | 1.148 | 0.56 | 58.67 | .579 |
| 4 | Group [OLZ3] | 1.195 | 1.148 | 1.04 | 58.67 | .302 |
| 5 | Group [OLZ6] | 1.493 | 1.148 | 1.30 | 58.67 | .198 |
| 6 | Phase [Post-Tx] | 1.884 | 0.930 | 2.03 | 48.43 | .048 |
| 7 | Phase [HF] | 11.960 | 0.934 | 12.81 | 49.23 | < .001 |
| 8 | PND*Group [No Dough] | 0.021 | 0.024 | 0.85 | 69.96 | .397 |
| 9 | PND*Group [OLZ3] | 0.053 | 0.024 | 2.19 | 69.96 | .032 |
| 10 | PND*Group [OLZ6] | 0.048 | 0.024 | 1.98 | 69.96 | .051 |
| 11 | PND*Phase [Post-Tx] | -0.042 | 0.022 | -1.89 | 56.11 | .064 |
| 12 | PND*Phase [HF] | 0.063 | 0.023 | 2.78 | 61.51 | .007 |
| 13 | Group [No Dough]*Phase [Post-Tx] | -0.009 | 1.315 | -0.01 | 48.43 | .995 |
| 14 | Group [OLZ3]*Phase [Post-Tx] | -0.269 | 1.315 | -0.20 | 48.43 | .839 |
| 15 | Group [OLZ6]*Phase [Post-Tx] | -0.148 | 1.315 | -0.11 | 48.43 | .911 |
| 16 | Group [No Dough]*Phase [HF] | -2.604 | 1.320 | -1.97 | 49.23 | .054 |
| 17 | Group [OLZ3]*Phase [HF] | -1.788 | 1.320 | -1.35 | 49.23 | .182 |
| 18 | Group [OLZ6]*Phase [HF] | 0.011 | 1.320 | 0.01 | 49.23 | .994 |
| 19 | PND*Group [No Dough]*Phase [Post-Tx] | -0.025 | 0.031 | -0.80 | 56.11 | .426 |
| 20 | PND*Group [OLZ3]*Phase [Post-Tx] | -0.055 | 0.031 | -1.76 | 56.11 | .083 |
| 21 | PND*Group [OLZ6]*Phase [Post-Tx] | -0.050 | 0.031 | -1.61 | 56.11 | .113 |
| 22 | PND*Group [No Dough]*Phase [HF] | -0.073 | 0.032 | -2.30 | 61.51 | .025 |
| 23 | PND*Group [OLZ3]*Phase [HF] | -0.109 | 0.032 | -3.41 | 61.51 | .001 |
| 24 | PND*Group [OLZ6]*Phase [HF] | -0.037 | 0.032 | -1.16 | 61.51 | .251 |
| | Combined | | | | | |
| 1 | (Intercept) | 17.841 | 0.382 | 46.73 | 162.89 | < .001 |
| 2 | PND | 0.068 | 0.008 | 8.34 | 194.66 | < .001 |
| 3 | Group [Treated] | 1.689 | 0.515 | 3.28 | 163.01 | .001 |
| 4 | Phase [Post-Tx] | 2.019 | 0.429 | 4.70 | 142.93 | < .001 |
| 5 | Phase [HF] | 10.368 | 0.432 | 24.00 | 146.63 | < .001 |
| 6 | PND*Group [Treated] | 0.048 | 0.011 | 4.34 | 195.19 | < .001 |
| 7 | PND*Phase [Post-Tx] | -0.035 | 0.011 | -3.31 | 167.93 | .001 |
| 8 | PND*Phase [HF] | 0.039 | 0.011 | 3.61 | 183.53 | < .001 |
| 9 | Group [Treated]*Phase [Post-Tx] | -0.493 | 0.579 | -0.85 | 143.01 | .396 |
| 10 | Group [Treated]*Phase [HF] | 0.529 | 0.582 | 0.91 | 146.68 | .365 |
| 11 | PND*Group [Treated]*Phase [Post-Tx] | -0.051 | 0.014 | -3.55 | 168.20 | < .001 |
| 12 | PND*Group [Treated]*Phase [HF] | -0.027 | 0.015 | -1.83 | 183.71 | .069 |

Abbreviations. Tx = Treatment Group (3 mg/kg/day risperidone in Risperidone-Treated Females Cohort 1; 0.5 mg/kg 2x/day risperidone in Risperidone-Treated Females Cohort 2; 3 mg/kg/day olanzapine in

Olanzapine-Treated Females Cohort 1 and 2); Post-Tx = Post-Treatment Phase; HF = High-Fat Diet Phase; No Dough = No Dough Control Group; Olz 3 = Olanzapine 3 mg/kg/day group; Olz 6 = Olanzapine 6 mg/kg/day.

Table S3. Regression parameter estimates from models of caloric intake.

Model term number, predictor, coefficient estimates (Estimate) of the predictor, standard errors (SE) of the estimates, and resulting t-statistic (t), degrees of freedom (df; Satterthwaite estimated), and p-values (p) from the multilevel linear regression analysis of caloric intake. All other details as described for Table S1.

| Model Term | Predictor | Estimate | SE | t | df | p |
|---|---|---|---|---|---|---|
| | EXP01: 3 mg/kg Risperidone | | | | | |
| 1 | (Intercept) | 7.118 | 0.451 | 15.79 | 173.02 | < .001 |
| 2 | PND | -0.133 | 0.018 | -7.28 | 1383.55 | < .001 |
| 3 | Group [Treated] | 1.544 | 0.639 | 2.42 | 174.60 | .017 |
| 4 | Phase [Post-Tx] | 1.725 | 0.475 | 3.63 | 121.70 | < .001 |
| 5 | Phase [HF] | 1.849 | 0.617 | 3.00 | 313.94 | .003 |
| 6 | PND*Group [Treated] | 0.042 | 0.026 | 1.63 | 1384.22 | .102 |
| 7 | PND*Phase [Post-Tx] | 0.124 | 0.019 | 6.59 | 1015.64 | < .001 |
| 8 | PND*Phase [HF] | 0.029 | 0.027 | 1.08 | 1421.85 | .282 |
| 9 | Group [Treated]*Phase [Post-Tx] | -1.408 | 0.674 | -2.09 | 123.31 | .039 |
| 10 | Group [Treated]*Phase [HF] | -1.619 | 0.873 | -1.85 | 315.31 | .065 |
| 11 | PND*Group [Treated]*Phase [Post-Tx] | -0.049 | 0.027 | -1.86 | 1016.47 | .064 |
| 12 | PND*Group [Treated]*Phase [HF] | -0.076 | 0.038 | -1.98 | 1422.15 | .047 |
| | EXP03: 0.5 mg/kg RIS 2x/daily | | | | | |
| 1 | (Intercept) | 10.517 | 0.788 | 13.35 | 41.44 | < .001 |
| 2 | PND | -0.047 | 0.028 | -1.70 | 389.08 | .090 |
| 3 | Group [Treated] | 2.963 | 1.114 | 2.66 | 41.44 | .011 |
| 4 | Phase [Post-Tx] | 0.309 | 0.751 | 0.41 | 31.22 | .684 |
| 5 | Phase [HF] | -0.347 | 0.897 | -0.39 | 62.81 | .700 |
| 6 | PND*Group [Treated] | 0.054 | 0.039 | 1.37 | 389.08 | .171 |
| 7 | PND*Phase [Post-Tx] | 0.051 | 0.030 | 1.74 | 191.11 | .083 |
| 8 | PND*Phase [HF] | -0.050 | 0.043 | -1.16 | 638.69 | .247 |
| 9 | Group [Treated]*Phase [Post-Tx] | -0.655 | 1.063 | -0.62 | 31.27 | .543 |
| 10 | Group [Treated]*Phase [HF] | -3.184 | 1.269 | -2.51 | 62.88 | .015 |
| 11 | PND*Group [Treated]*Phase [Post-Tx] | -0.059 | 0.042 | -1.41 | 191.32 | .160 |
| 12 | PND*Group [Treated]*Phase [HF] | -0.170 | 0.061 | -2.80 | 642.23 | .005 |
| | EXP02: 3 mg/kg Olanzapine | | | | | |
| 1 | (Intercept) | 8.220 | 0.372 | 22.13 | 102.93 | < .001 |
| 2 | PND | -0.079 | 0.013 | -6.19 | 288.41 | < .001 |
| 3 | Group [Treated] | 1.389 | 0.525 | 2.64 | 102.93 | .009 |
| 4 | Phase [Post-Tx] | 1.340 | 0.376 | 3.57 | 84.16 | < .001 |
| 5 | Phase [HF] | -0.065 | 0.913 | -0.07 | 1302.40 | .944 |
| 6 | PND*Group [Treated] | 0.024 | 0.018 | 1.34 | 288.63 | .180 |
| 7 | PND*Phase [Post-Tx] | 0.089 | 0.013 | 6.92 | 199.53 | < .001 |
| 8 | PND*Phase [HF] | -0.037 | 0.030 | -1.24 | 1613.00 | .215 |
| 9 | Group [Treated]*Phase [Post-Tx] | -1.227 | 0.531 | -2.31 | 84.10 | .023 |
| 10 | Group [Treated]*Phase [HF] | -0.537 | 1.287 | -0.42 | 1295.72 | .677 |

| | | | | | | |
|---|---|---|---|---|---|---|
| 11 | PND*Group [Treated]*Phase [Post-Tx] | -0.023 | 0.018 | -1.25 | 199.81 | .213 |
| 12 | PND*Group [Treated]*Phase [HF] | -0.017 | 0.042 | -0.39 | 1609.45 | .693 |
| | EXP04: 3 and 6 mg/kg Olanzapine | | | | | |
| 1 | (Intercept) | 11.354 | 0.664 | 17.09 | 134.92 | < .001 |
| 2 | PND | -0.025 | 0.024 | -1.00 | 2429.92 | .316 |
| 3 | Group [No Dough] | 4.062 | 0.950 | 4.27 | 140.94 | < .001 |
| 4 | Group [OLZ3] | 2.863 | 0.947 | 3.02 | 139.30 | .003 |
| 5 | Group [OLZ6] | -0.044 | 0.940 | -0.05 | 134.92 | .962 |
| 6 | Phase [Post-Tx] | -0.093 | 0.696 | -0.13 | 93.80 | .895 |
| 7 | Phase [HF] | -1.635 | 0.842 | -1.94 | 197.39 | .054 |
| 8 | PND*Group [No Dough] | 0.118 | 0.035 | 3.36 | 2458.27 | < .001 |
| 9 | PND*Group [OLZ3] | 0.070 | 0.035 | 2.00 | 2448.99 | .046 |
| 10 | PND*Group [OLZ6] | -0.019 | 0.035 | -0.55 | 2429.92 | .584 |
| 11 | PND*Phase [Post-Tx] | 0.026 | 0.025 | 1.03 | 1481.30 | .302 |
| 12 | PND*Phase [HF] | -0.114 | 0.037 | -3.11 | 2425.28 | .002 |
| 13 | Group [No Dough]*Phase [Post-Tx] | -1.026 | 0.995 | -1.03 | 97.93 | .305 |
| 14 | Group [OLZ3]*Phase [Post-Tx] | -1.880 | 0.992 | -1.89 | 96.78 | .061 |
| 15 | Group [OLZ6]*Phase [Post-Tx] | -0.254 | 0.984 | -0.26 | 93.80 | .797 |
| 16 | Group [No Dough]*Phase [HF] | -4.274 | 1.199 | -3.56 | 202.61 | < .001 |
| 17 | Group [OLZ3]*Phase [HF] | -3.205 | 1.197 | -2.68 | 201.24 | .008 |
| 18 | Group [OLZ6]*Phase [HF] | 0.173 | 1.192 | 0.15 | 198.21 | .885 |
| 19 | PND*Group [No Dough]*Phase [Post-Tx] | -0.152 | 0.036 | -4.24 | 1531.53 | < .001 |
| 20 | PND*Group [OLZ3]*Phase [Post-Tx] | -0.091 | 0.036 | -2.54 | 1517.21 | .011 |
| 21 | PND*Group [OLZ6]*Phase [Post-Tx] | 0.014 | 0.035 | 0.39 | 1481.37 | .697 |
| 22 | PND*Group [No Dough]*Phase [HF] | -0.113 | 0.052 | -2.16 | 2438.20 | .031 |
| 23 | PND*Group [OLZ3]*Phase [HF] | -0.101 | 0.052 | -1.93 | 2433.75 | .054 |
| 24 | PND*Group [OLZ6]*Phase [HF] | 0.010 | 0.052 | 0.18 | 2430.18 | .855 |
| | Combined | | | | | |
| 1 | (Intercept) | 9.233 | 0.336 | 27.44 | 195.57 | < .001 |
| 2 | PND | -0.077 | 0.010 | -7.99 | 1556.42 | < .001 |
| 3 | Group [Treated] | 2.021 | 0.456 | 4.43 | 200.67 | < .001 |
| 4 | Phase [Post-Tx] | 0.841 | 0.310 | 2.71 | 132.81 | .008 |
| 5 | Phase [HF] | 0.270 | 0.390 | 0.69 | 323.12 | .489 |
| 6 | PND*Group [Treated] | 0.037 | 0.013 | 2.85 | 1624.88 | .004 |
| 7 | PND*Phase [Post-Tx] | 0.079 | 0.010 | 7.76 | 305.77 | < .001 |
| 8 | PND*Phase [HF] | -0.026 | 0.015 | -1.68 | 1394.48 | .094 |
| 9 | Group [Treated]*Phase [Post-Tx] | -1.141 | 0.421 | -2.71 | 136.76 | .008 |
| 10 | Group [Treated]*Phase [HF] | -1.964 | 0.529 | -3.71 | 331.47 | < .001 |
| 11 | PND*Group [Treated]*Phase [Post-Tx] | -0.045 | 0.014 | -3.20 | 320.76 | .001 |
| 12 | PND*Group [Treated]*Phase [HF] | -0.074 | 0.021 | -3.50 | 1433.31 | < .001 |

Abbreviations as described for Table S2.

Table S4. Intercepts, coefficients of age and between-group contrasts of estimates from body weight models.

Estimates for the comparison groups ("Est. 1") and the plain dough control group ("Est. 2") along with standard errors (SE) and within-phase contrasts versus the plain dough control group of intercepts (starting body weight) and slope (change in body weight per day). The emmeans command was used to compare intercept estimates and the emtrends command was used to compare slopes and resulting t-statistic values [degrees of freedom] and associated p-values for each contrast are provided in the Statistical Comparison column. P-values were Tukey-adjusted for multiple comparisons.

| **Experiment** | **Phase** | **Contrast** | **Est. 1** | **Est. 2** | **Diff.** | **SE** | **t** | **df** | **p** |
|---|---|---|---|---|---|---|---|---|---|
| | | | *Intercept* | | | | | | |
| EXP01: 3 mg/kg Risperidone | Tx | Treatment - Control | 15.5 | 15.5 | 0.0 | 0.504 | 0.10 | 26.94 | .923 |
| EXP01: 3 mg/kg Risperidone | Post-Tx | Treatment - Control | 20.7 | 19.3 | 1.4 | 0.502 | 2.79 | 26.51 | .010 |
| EXP01: 3 mg/kg Risperidone | HF | Treatment - Control | 25.3 | 23.9 | 1.5 | 0.504 | 2.94 | 26.94 | .007 |
| EXP03: 0.5 mg/kg RIS 2x/daily | Tx | Treatment - Control | 16.1 | 15.4 | 0.7 | 0.561 | 1.30 | 34.58 | .202 |
| EXP03: 0.5 mg/kg RIS 2x/daily | Post-Tx | Treatment - Control | 19.8 | 17.9 | 1.8 | 0.555 | 3.32 | 33.19 | .002 |
| EXP03: 0.5 mg/kg RIS 2x/daily | HF | Treatment - Control | 26.2 | 24.0 | 2.2 | 0.562 | 4.00 | 34.76 | < .001 |
| EXP02: 3 mg/kg Olanzapine | Tx | Treatment - Control | 15.6 | 15.8 | -0.2 | 0.609 | -0.25 | 25.61 | .801 |
| EXP02: 3 mg/kg Olanzapine | Post-Tx | Treatment - Control | 19.9 | 19.0 | 0.8 | 0.607 | 1.36 | 25.21 | .186 |
| EXP02: 3 mg/kg Olanzapine | HF | Treatment - Control | 25.5 | 24.9 | 0.6 | 0.613 | 1.00 | 26.22 | .328 |
| EXP04: 3 and 6 mg/kg Olanzapine | Tx | NoDough - Control | 15.6 | 15.6 | 0.0 | 0.702 | 0.03 | 59.09 | 1.000 |
| EXP04: 3 and 6 mg/kg Olanzapine | Tx | OLZ3 - Control | 15.2 | 15.6 | -0.4 | 0.702 | -0.56 | 59.09 | .871 |
| EXP04: 3 and 6 mg/kg Olanzapine | Tx | OLZ6 - Control | 15.6 | 15.6 | 0.1 | 0.702 | 0.08 | 59.09 | .997 |
| EXP04: 3 and 6 mg/kg Olanzapine | Post-Tx | NoDough - Control | 19.5 | 18.7 | 0.8 | 0.696 | 1.10 | 57.29 | .555 |
| EXP04: 3 and 6 mg/kg Olanzapine | Post-Tx | OLZ3 - Control | 19.7 | 18.7 | 1.0 | 0.696 | 1.42 | 57.29 | .362 |
| EXP04: 3 and 6 mg/kg Olanzapine | Post-Tx | OLZ6 - Control | 20.1 | 18.7 | 1.4 | 0.696 | 2.03 | 57.29 | .122 |
| EXP04: 3 and 6 mg/kg Olanzapine | HF | NoDough - Control | 25.3 | 25.6 | -0.4 | 0.702 | -0.55 | 59.01 | .875 |
| EXP04: 3 and 6 mg/kg Olanzapine | HF | OLZ3 - Control | 26.7 | 25.6 | 1.1 | 0.702 | 1.55 | 59.01 | .296 |
| EXP04: 3 and 6 mg/kg Olanzapine | HF | OLZ6 - Control | 26.8 | 25.6 | 1.2 | 0.702 | 1.68 | 59.01 | .238 |
| Combined | Tx | Treatment - Control | 15.6 | 15.5 | 0.1 | 0.293 | 0.26 | 169.87 | .795 |
| Combined | Post-Tx | Treatment - Control | 20.0 | 18.7 | 1.3 | 0.291 | 4.43 | 165.83 | < .001 |
| Combined | HF | Treatment - Control | 26.1 | 24.6 | 1.5 | 0.293 | 5.12 | 170.62 | < .001 |

| | | | | | | | | | |
|---|---|---|---|---|---|---|---|---|---|
| | | | *Slope* | | | | | | |
| EXP01: 3 mg/kg Risperidone | Tx | Treatment - Control | 0.13 | 0.08 | 0.04 | 0.024 | 1.86 | 39.50 | .070 |
| EXP01: 3 mg/kg Risperidone | Post-Tx | Treatment - Control | 0.01 | 0.02 | -0.01 | 0.023 | -0.42 | 35.75 | .675 |
| EXP01: 3 mg/kg Risperidone | HF | Treatment - Control | 0.13 | 0.12 | 0.01 | 0.024 | 0.53 | 39.88 | .596 |
| EXP03: 0.5 mg/kg RIS 2x/daily | Tx | Treatment - Control | 0.10 | 0.05 | 0.05 | 0.022 | 2.49 | 43.39 | .017 |
| EXP03: 0.5 mg/kg RIS 2x/daily | Post-Tx | Treatment - Control | 0.03 | 0.04 | -0.01 | 0.021 | -0.36 | 36.12 | .724 |
| EXP03: 0.5 mg/kg RIS 2x/daily | HF | Treatment - Control | 0.15 | 0.11 | 0.04 | 0.022 | 1.95 | 45.91 | .058 |
| EXP02: 3 mg/kg Olanzapine | Tx | Treatment - Control | 0.09 | 0.06 | 0.03 | 0.023 | 1.27 | 40.29 | .211 |
| EXP02: 3 mg/kg Olanzapine | Post-Tx | Treatment - Control | 0.03 | 0.03 | 0.00 | 0.023 | 0.05 | 38.58 | .963 |
| EXP02: 3 mg/kg Olanzapine | HF | Treatment - Control | 0.12 | 0.08 | 0.04 | 0.024 | 1.65 | 47.52 | .105 |
| EXP04: 3 and 6 mg/kg Olanzapine | Tx | NoDough - Control | 0.10 | 0.08 | 0.02 | 0.024 | 0.85 | 74.08 | .708 |
| EXP04: 3 and 6 mg/kg Olanzapine | Tx | OLZ3 - Control | 0.14 | 0.08 | 0.05 | 0.024 | 2.19 | 74.08 | .084 |
| EXP04: 3 and 6 mg/kg Olanzapine | Tx | OLZ6 - Control | 0.13 | 0.08 | 0.05 | 0.024 | 1.98 | 74.08 | .131 |
| EXP04: 3 and 6 mg/kg Olanzapine | Post-Tx | NoDough - Control | 0.04 | 0.04 | 0.00 | 0.023 | -0.19 | 62.63 | .986 |
| EXP04: 3 and 6 mg/kg Olanzapine | Post-Tx | OLZ3 - Control | 0.04 | 0.04 | 0.00 | 0.023 | -0.09 | 62.63 | .997 |
| EXP04: 3 and 6 mg/kg Olanzapine | Post-Tx | OLZ6 - Control | 0.04 | 0.04 | 0.00 | 0.023 | -0.10 | 62.64 | .996 |
| EXP04: 3 and 6 mg/kg Olanzapine | HF | NoDough - Control | 0.09 | 0.15 | -0.05 | 0.024 | -2.18 | 73.75 | .085 |
| EXP04: 3 and 6 mg/kg Olanzapine | HF | OLZ3 - Control | 0.09 | 0.15 | -0.06 | 0.024 | -2.32 | 73.75 | .062 |
| EXP04: 3 and 6 mg/kg Olanzapine | HF | OLZ6 - Control | 0.16 | 0.15 | 0.01 | 0.024 | 0.45 | 73.75 | .914 |
| Combined | Tx | Treatment - Control | 0.12 | 0.07 | 0.05 | 0.011 | 4.28 | 211.99 | < .001 |
| Combined | Post-Tx | Treatment - Control | 0.03 | 0.03 | 0.00 | 0.011 | -0.26 | 189.84 | .798 |
| Combined | HF | Treatment - Control | 0.13 | 0.11 | 0.02 | 0.011 | 1.88 | 219.42 | .062 |

Table S5. Intercepts, coefficients of age and between-group contrasts of estimates from caloric intake models.

Within-cohort and within-phase contrasts versus the control group of intercepts (starting caloric intake). Estimate of the intercept for the first group in the comparison ("Comparator") and the control group ("Control") along with the standard error (SE) of the estimate are provided for the intercept of the model along with the estimated difference and its SE. The emmeans command was used to conduct contrasts and resulting t-statistic values [degrees of freedom] and associated p-values provided in the Statistical Comparison column. P-values were Tukey-adjusted for multiple comparisons in the third olanzapine cohort.

| **Experiment** | **Phase** | **Contrast** | **Est. 1** | **Est. 2** | **Diff.** | **SE** | **t** | **df** | **p** |
|---|---|---|---|---|---|---|---|---|---|
| | | | *Intercept* | | | | | | |
| EXP01: 3 mg/kg Risperidone | Tx | Treatment - Control | 11.7 | 11.5 | 0.1 | 0.614 | 0.23 | 61.97 | .821 |
| EXP01: 3 mg/kg Risperidone | Post-Tx | Treatment - Control | 9.5 | 9.1 | 0.4 | 0.575 | 0.66 | 47.73 | .513 |
| EXP01: 3 mg/kg Risperidone | HF | Treatment - Control | 13.5 | 12.4 | 1.1 | 0.636 | 1.67 | 71.01 | .099 |
| EXP03: 0.5 mg/kg RIS 2x/daily | Tx | Treatment - Control | 13.3 | 11.8 | 1.5 | 1.106 | 1.35 | 44.44 | .185 |
| EXP03: 0.5 mg/kg RIS 2x/daily | Post-Tx | Treatment - Control | 13.2 | 10.7 | 2.4 | 1.011 | 2.42 | 31.05 | .022 |
| EXP03: 0.5 mg/kg RIS 2x/daily | HF | Treatment - Control | 15.8 | 12.8 | 3.0 | 1.148 | 2.59 | 51.53 | .012 |
| EXP02: 3 mg/kg Olanzapine | Tx | Treatment - Control | 11.9 | 11.5 | 0.4 | 0.697 | 0.53 | 32.28 | .596 |
| EXP02: 3 mg/kg Olanzapine | Post-Tx | Treatment - Control | 9.2 | 9.1 | 0.1 | 0.651 | 0.16 | 24.51 | .877 |
| EXP02: 3 mg/kg Olanzapine | HF | Treatment - Control | 13.6 | 13.0 | 0.5 | 0.792 | 0.68 | 53.33 | .500 |
| EXP04: 3 and 6 mg/kg Olanzapine | Tx | NoDough - Control | 12.6 | 12.1 | 0.5 | 0.965 | 0.54 | 81.17 | .880 |
| EXP04: 3 and 6 mg/kg Olanzapine | Tx | OLZ3 - Control | 12.9 | 12.1 | 0.8 | 0.965 | 0.79 | 81.26 | .746 |
| EXP04: 3 and 6 mg/kg Olanzapine | Tx | OLZ6 - Control | 12.6 | 12.1 | 0.5 | 0.963 | 0.55 | 80.79 | .876 |
| EXP04: 3 and 6 mg/kg Olanzapine | Post-Tx | NoDough - Control | 15.3 | 11.2 | 4.1 | 0.866 | 4.69 | 52.74 | < .001 |
| EXP04: 3 and 6 mg/kg Olanzapine | Post-Tx | OLZ3 - Control | 12.8 | 11.2 | 1.6 | 0.865 | 1.85 | 52.63 | .173 |
| EXP04: 3 and 6 mg/kg Olanzapine | Post-Tx | OLZ6 - Control | 11.1 | 11.2 | -0.1 | 0.863 | -0.16 | 52.17 | .989 |
| EXP04: 3 and 6 mg/kg Olanzapine | HF | NoDough - Control | 13.5 | 13.9 | -0.4 | 0.991 | -0.38 | 90.30 | .941 |
| EXP04: 3 and 6 mg/kg Olanzapine | HF | OLZ3 - Control | 14.5 | 13.9 | 0.6 | 0.991 | 0.58 | 90.30 | .858 |
| EXP04: 3 and 6 mg/kg Olanzapine | HF | OLZ6 - Control | 14.3 | 13.9 | 0.4 | 0.995 | 0.41 | 91.62 | .929 |
| Combined | Tx | Treatment - Control | 12.6 | 11.8 | 0.8 | 0.457 | 1.66 | 194.36 | .098 |
| Combined | Post-Tx | Treatment - Control | 11.1 | 10.0 | 1.1 | 0.428 | 2.62 | 148.43 | .010 |
| Combined | HF | Treatment - Control | 14.2 | 13.0 | 1.3 | 0.475 | 2.69 | 227.49 | .008 |
| | | | *Slope* | | | | | | |

| | | | | | | | | | |
|---|---|---|---|---|---|---|---|---|---|
| EXP01: 3 mg/kg Risperidone | Tx | Treatment - Control | -0.09 | -0.13 | 0.04 | 0.026 | 1.63 | 1266.63 | .102 |
| EXP01: 3 mg/kg Risperidone | Post-Tx | Treatment - Control | -0.02 | -0.01 | -0.01 | 0.007 | -1.10 | 19.83 | .284 |
| EXP01: 3 mg/kg Risperidone | HF | Treatment - Control | -0.14 | -0.10 | -0.03 | 0.029 | -1.20 | 1372.02 | .231 |
| EXP03: 0.5 mg/kg RIS 2x/daily | Tx | Treatment - Control | 0.01 | -0.05 | 0.05 | 0.039 | 1.37 | 479.90 | .171 |
| EXP03: 0.5 mg/kg RIS 2x/daily | Post-Tx | Treatment - Control | 0.00 | 0.00 | -0.01 | 0.016 | -0.32 | 15.13 | .750 |
| EXP03: 0.5 mg/kg RIS 2x/daily | HF | Treatment - Control | -0.21 | -0.10 | -0.12 | 0.047 | -2.48 | 770.92 | .013 |
| EXP02: 3 mg/kg Olanzapine | Tx | Treatment - Control | -0.05 | -0.08 | 0.02 | 0.018 | 1.34 | 308.31 | .180 |
| EXP02: 3 mg/kg Olanzapine | Post-Tx | Treatment - Control | 0.01 | 0.01 | 0.00 | 0.009 | 0.16 | 18.39 | .871 |
| EXP02: 3 mg/kg Olanzapine | HF | Treatment - Control | -0.11 | -0.12 | 0.01 | 0.039 | 0.19 | 1688.91 | .848 |
| EXP04: 3 and 6 mg/kg Olanzapine | Tx | NoDough - Control | 0.09 | -0.02 | 0.12 | 0.035 | 3.36 | 2396.53 | .002 |
| EXP04: 3 and 6 mg/kg Olanzapine | Tx | OLZ3 - Control | 0.05 | -0.02 | 0.07 | 0.035 | 2.00 | 2386.13 | .119 |
| EXP04: 3 and 6 mg/kg Olanzapine | Tx | OLZ6 - Control | -0.04 | -0.02 | -0.02 | 0.035 | -0.55 | 2364.87 | .874 |
| EXP04: 3 and 6 mg/kg Olanzapine | Post-Tx | NoDough - Control | -0.03 | 0.00 | -0.03 | 0.008 | -4.36 | 26.52 | < .001 |
| EXP04: 3 and 6 mg/kg Olanzapine | Post-Tx | OLZ3 - Control | -0.02 | 0.00 | -0.02 | 0.008 | -2.63 | 26.56 | .038 |
| EXP04: 3 and 6 mg/kg Olanzapine | Post-Tx | OLZ6 - Control | 0.00 | 0.00 | -0.01 | 0.008 | -0.67 | 25.50 | .814 |
| EXP04: 3 and 6 mg/kg Olanzapine | HF | NoDough - Control | -0.13 | -0.14 | 0.01 | 0.039 | 0.14 | 2532.22 | .992 |
| EXP04: 3 and 6 mg/kg Olanzapine | HF | OLZ3 - Control | -0.17 | -0.14 | -0.03 | 0.039 | -0.79 | 2532.22 | .744 |
| EXP04: 3 and 6 mg/kg Olanzapine | HF | OLZ6 - Control | -0.15 | -0.14 | -0.01 | 0.039 | -0.24 | 2538.87 | .976 |
| Combined | Tx | Treatment - Control | -0.04 | -0.08 | 0.04 | 0.013 | 2.83 | 1983.26 | .005 |
| Combined | Post-Tx | Treatment - Control | 0.00 | 0.00 | -0.01 | 0.005 | -1.38 | 79.50 | .171 |
| Combined | HF | Treatment - Control | -0.14 | -0.10 | -0.04 | 0.017 | -2.18 | 3949.55 | .029 |

Note. Abbreviations as described for Table S3.

Figure S1. Body weight and caloric intake in male mice treated with 3 mg/kg risperidone.

A) body weight (g) in male C57BL/6J mice treated with 3 mg/kg/day risperidone in cookie dough or given plain dough from PND 37 – 66 ("Tx"), following risperidone or plain dough self administration from PND 67-196 ("Post-Tx") and during a period of high-fat diet feeding from PND 197-229 ("HF"). B) caloric intake (kcal) in the same mice across the treatment, post-treatment, and high-fat diet feeding phases. Data points represent group averages. Solid lines represent best-fitting regression model. Shaded areas around lines indicate the 95% confidence limits around the model fit.

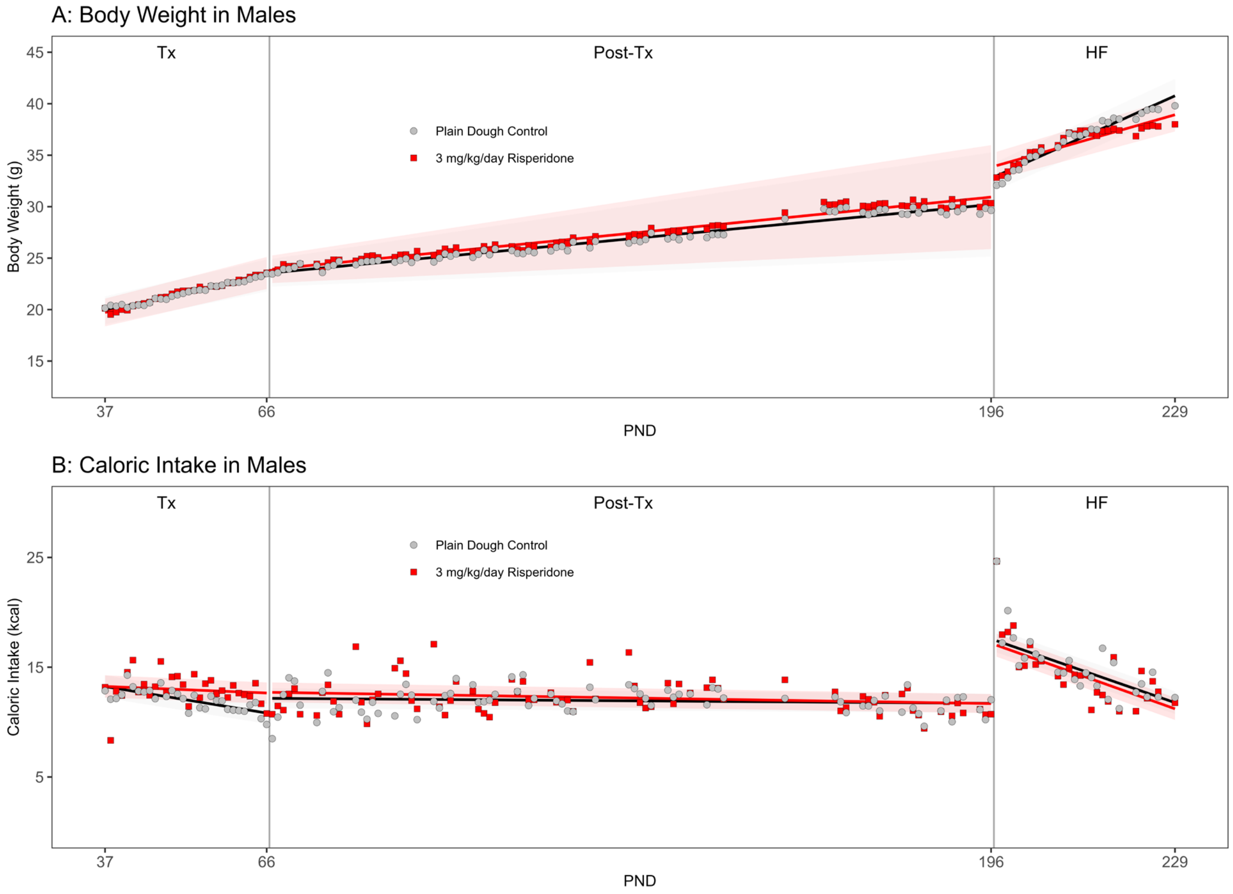

Figure S2. Circulating insulin and glucose.

Concentration of insulin in serum and glucose in blood following a 5-hour fast for mice previously treated with risperidone or olanzapine. **A**: Serum insulin concentration (check units) in second cohort of risperidone-treated female mice. **B**: Serum insulin concentration in third cohort of olanzapine-treated female mice. **C**: Blood glucose concentration in second cohort of risperidone-treated female mice. **D**: Blood glucose concentration in third cohort of olanzapine-treated female mice. Black horizontal bars represent group mean. Each individual data point represents result from an individual mouse.

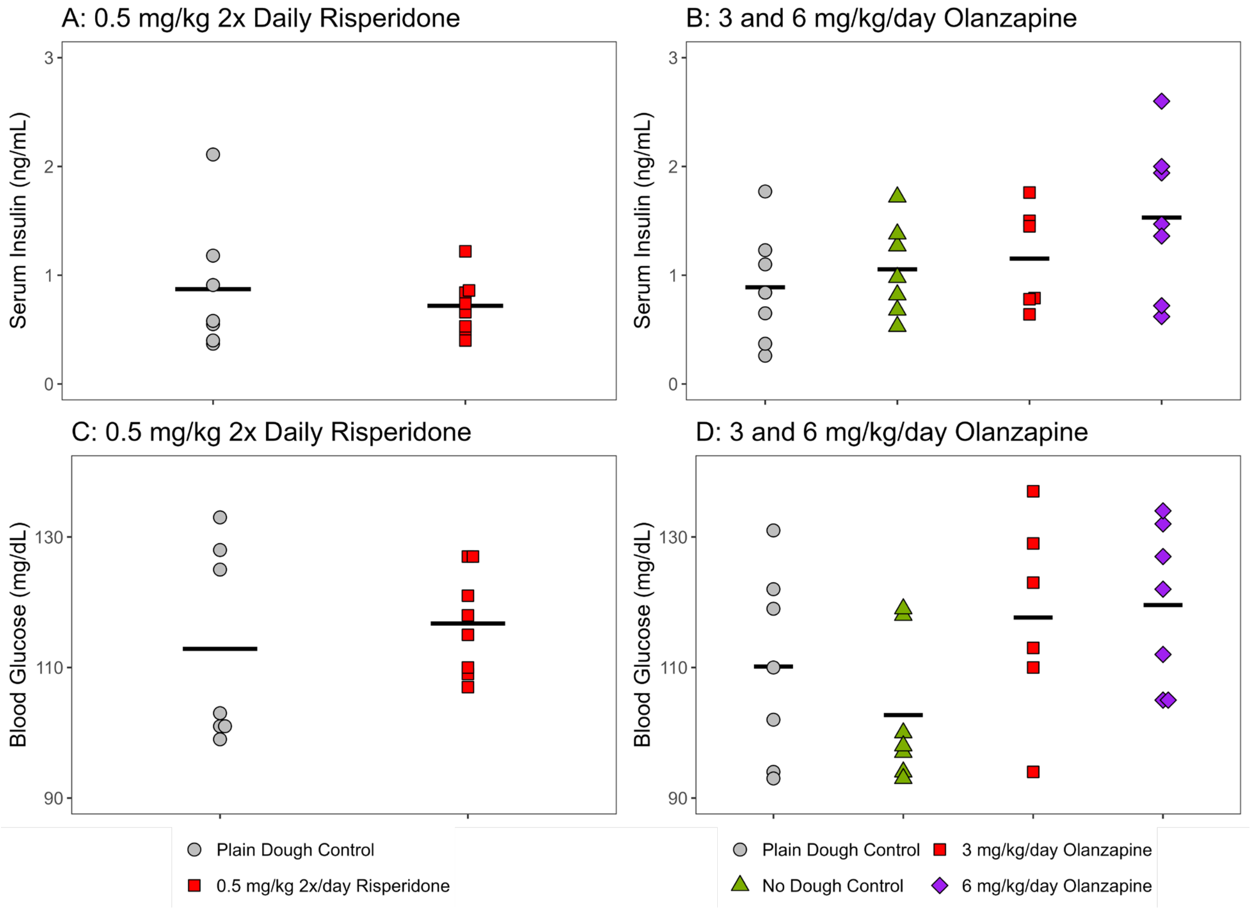
